# *SSARP*: An R Package for Easily Creating Species- and Speciation- Area Relationships Using Web Databases

**DOI:** 10.1101/2024.12.31.630948

**Authors:** Kristen M. Martinet, Cristian Román-Palacios, Luke J. Harmon

## Abstract

A universal method of quantifying patterns of biodiversity on islands is the species-area relationship (SAR). SARs visualize the relationship between species richness (the number of species) and the area of the land mass on which they live. An extension of this visualization, the speciation-area relationship (SpAR), helps researchers determine trends in speciation rate over a set of land masses. Comparing these relationships across island systems globally is an extremely difficult task because gathering and processing a large amount of species occurrence data and island data often requires researchers to conduct lengthy literature searches and combine datasets from several different sources. Here we present *SSARP* (Species/Speciation-Area Relationship Projector), an R package that provides a simple workflow for creating SARs and SpARs. The *SSARP* workflow allows users to gather occurrence data from GBIF, use mapping tools to determine whether the GPS points in the occurrence data refer to valid land masses, associate those land masses with their areas using a built-in dataset of island names and areas, and create SARs using linear and segmented regression. *SSARP* also provides multiple functions for estimating speciation rates for use in creating a SpAR. Using *SSARP* allows researchers to dramatically increase the scope of their biodiversity research through the creation of SARs and SpARs with data from island systems across the globe.

## Introduction

MacArthur and Wilson’s (1967) equilibrium model of island biogeography introduced a foundational framework for understanding patterns of biodiversity on islands. This original framework proposed an equilibrium point between immigration and extinction rates, and that the dynamics of this relationship would change based on island size and isolation from other land masses. This model of island biogeography has been expanded upon multiple times, with important additions relating to phylogenetic diversification (Heaney 2000), island ontogeny (Whittaker et al. 2008), and species abundances (Rosindell and Harmon 2013). One important way to help quantify patterns of biodiversity on islands within these frameworks is the species-area relationship (SAR), which visualizes the relationship between species richness (the number of species) and the area of the island on which they live. Additionally, the potential for speciation on islands can be visualized using a speciation-area relationship (SpAR), which plots speciation rates against the area of the island on which the associated species live.

The general observation that species richness increases with increasing area is a fundamental law of ecology (Arrhenius 1921; Gleason 1922; Rosenzweig 1995). Disruption of this relationship may be associated with decreasing biodiversity due to habitat loss and fragmentation (Chisholm et al. 2018) and increasing numbers of non-native species (Basier and Li 2018; Guo et al. 2021). Creating SARs for island-dwelling species helps researchers understand how trends in biodiversity across archipelagos are changing due to these effects. The trends in species richness that are visualized through the use of SARs can be further explained through the use of SpARs. SpARs help pinpoint a threshold land area for *in situ* speciation (Losos and Schluter 2000) or, in contrast, find that *in situ* speciation occurs regardless of area (Wagner et al. 2014).

Global comparisons of island systems are not common in the island biogeography literature because of the difficulty associated with gathering and processing the large amount of data needed to make informed conclusions. Researchers who have tackled this problem typically conduct a lengthy literature search and combine datasets from as many SAR-related papers as possible (e.g. Baiser and Li 2018; Guo et al. 2021; da Silva et al. 2024). Given the increasing availability of occurrence data from databases such as GBIF (Global Biodiversity Information Facility), creating SARs to visualize patterns of biodiversity on a global scale should be a more accessible practice. Our R package *SSARP* (Species/Speciation-Area Relationship Projector) streamlines the process of using global biodiversity data from GBIF to create species-area and speciation-area relationships.

SARs require two main components to be constructed: occurrence data for taxa of interest and the area for each land mass on which those taxa reside. SpARs need these two components, along with a phylogeny to use in estimating speciation rates. A popular method of compiling occurrence data is through accessing GBIF and other web databases of occurrence data via their APIs using R packages such as *rgbif* (Chamberlain et al. 2024) or *spocc* (Owens et al. 2024). There are several options for filtering data using these packages, such as including only occurrence data with GPS points or restricting the geographical area of the data it returns, but sometimes this data still requires manual cleaning. The *CoordinateCleaner* R package (Zizka et al. 2019) provides a suite of useful tools for ensuring that occurrence data from web databases has meaningful GPS points for use in analyses (for example, testing if GPS points plot in the ocean instead of on land, if locality information matches the true location of the GPS points, and if the GPS points correspond with museum locations instead of true observations). However, the tools available with *CoordinateCleaner* frequently flag valid occurrence records on small islands when a buffer for the coastline is not implemented, and implementing this kind of buffer is often tedious for islands with minimal geographic data available on global datasets. *SSARP* fixes this flagging problem by combining different sources of geographic information to determine whether occurrence points are truly on land.

*SSARP* uses the utility of *rgbif* (Chamberlain et al. 2024) to access occurrence records on GBIF, which is the occurrence database of choice for this R package because it includes data from several different online occurrence record databases. To help with the creation of SARs and SpARs, especially for island-dwelling species, *SSARP* also includes a built-in dataset of island names and their associated areas that was created using global island data from Sayre et al. (2018) with ArcGIS Pro (ESRI 2024). Once occurrence data has been curated and the land mass area is recorded, the next step for creating these relationships is to use a regression model to describe the relationship between the number of species (or speciation rate in the case of SpARs) and the area of the land mass on which they live. For island systems, this regression model is often represented by a linear regression on a log-log scale (reviewed in Scheiner 2003) or a segmented regression (reviewed in Matthews and Rigal 2021). *SSARP* provides functionality that allows users to create both linear regression models and segmented regression models. However, we acknowledge that there are several alternative models that can be used to fit SARs. For instance, the R package *sars* (Matthews et al. 2019) includes functions to fit SARs using 20 different models and includes methodology for several SAR-related analyses. *SSARP* includes only basic functions for fitting SARs because we primarily focus on accessing data, filtering out invalid occurrence records, and providing speciation rate methodology for creating SpARs.

## Description

The *SSARP* package allows users to create species-area relationships (SARs) for a taxon of interest (e.g., the name of a species, subspecies, genus, family) through the use of functions that serve as sequential steps in the workflow of creating a SAR (Table 1).

**Table 1.** Functions included in *SSARP* for creating a species-area relationship (SAR).

| Function | Description | Input |
| --- | --- | --- |
| getKey | Use *rgbif* (Chamberlain et al. 2024) to find the best match for the name given and return the taxon key | Taxon name and rank. |
| getData | Use *rgbif* (Chamberlain et al. 2024) to retrieve occurrence data from GBIF's database for a given taxon. This function will only return occurrences that include GPS coordinates. | Taxon key from "getKey," record number limit, (optional) geographic constraint in Well Known Text (WKT) format. |
| findLand | Use various mapping tools to attempt to find the names of land masses where the occurrence points were found. | Occurrence records, whether the function should use the Photon API to double-check points that were categorized as not on land (TRUE) or not (FALSE). |
| findAreas | Reference a dataset of island names and areas to find the areas of the land masses relevant to the taxon of interest. | Occurrence records from "findLand" |
| removeContinents | Reference a list of continental areas to remove them from the dataframe output by the findAreas function. | Occurrence records from "findAreas" |
| SARP | Use linear or segmented regression up to a specified number of breakpoints to create species-area relationship plots (SARP). The best model as determined by AIC is returned. | Occurrence records from "findAreas," maximum number of breakpoints to include in regression model selection. |
| quickSARP | Use the basic SSARP workflow for creating a species-area relationship for island species using occurrence data from GBIF without having to execute every step manually. | Taxon name, taxon rank, occurrence record number limit, (optional) geographic constraint in WKT format, maximum number of breakpoints to include in SAR. |

The basic workflow for *SSARP* when the user wants to create an SAR involves the following steps: (1) gather data from GBIF, (2) determine whether the GPS points associated with the occurrence records truly correspond with land masses, (3) find the areas of those landmasses from a dataset included in the package, and (4) create a species-area relationship using the resulting curated data (Figure 1).

**Figure 1.**
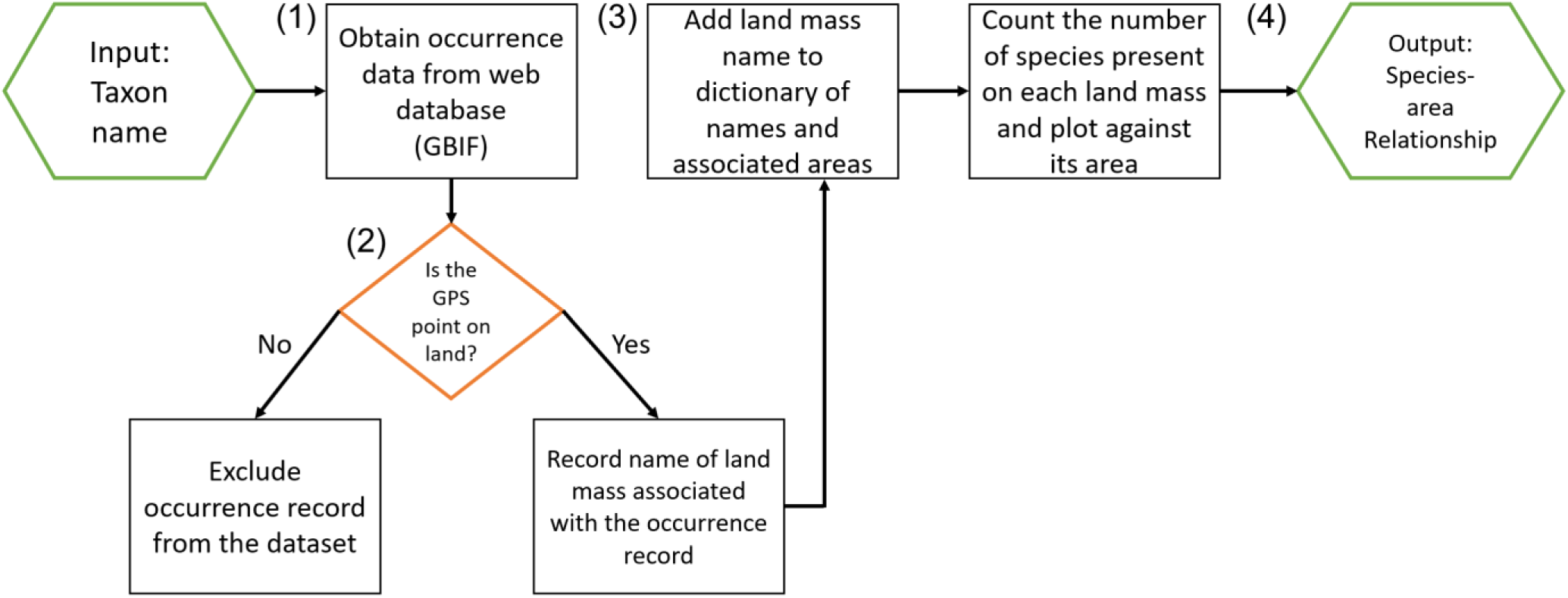
A flowchart representing the basic workflow for using *SSARP* to create a species-area relationship. Steps for creating the SAR are numbered: (1) gather data from GBIF, (2) determine whether the GPS points associated with the occurrence records truly correspond with land masses, (3) find the areas of those landmasses from a dataset included in the package, and (4) create a species-area relationship using the resulting curated data.

To create a SpAR for a taxon of interest, the user will follow the same workflow as with SARs, but with an extra user-specified method for estimating speciation rates (Figure 2). Once a dataset including occurrence records on land and the areas of their associated landmasses is created using the *SSARP* workflow, the user must provide a phylogenetic tree for the taxa of interest and select a method for estimating speciation rates. *SSARP* currently supports three different methods for estimating speciation rates: BAMM (Rabosky 2014), the lambda calculation for crown groups from Magallόn and Sanderson (2001), and DR (Jetz et al. 2012). In order to use the BAMM method for estimating speciation rates, the user must supply a bammdata object generated by reading the event data file from a BAMM analysis with the *BAMMtools* package (Rabosky et al. 2014). In order to use the Magallόn and Sanderson (2001) and the DR (Jetz et al. 2012) methods, the user must input a phylogenetic tree associated with the taxa of interest. Functions specifically relevant to creating SpARs are described in Table 2.

**Figure 2.**
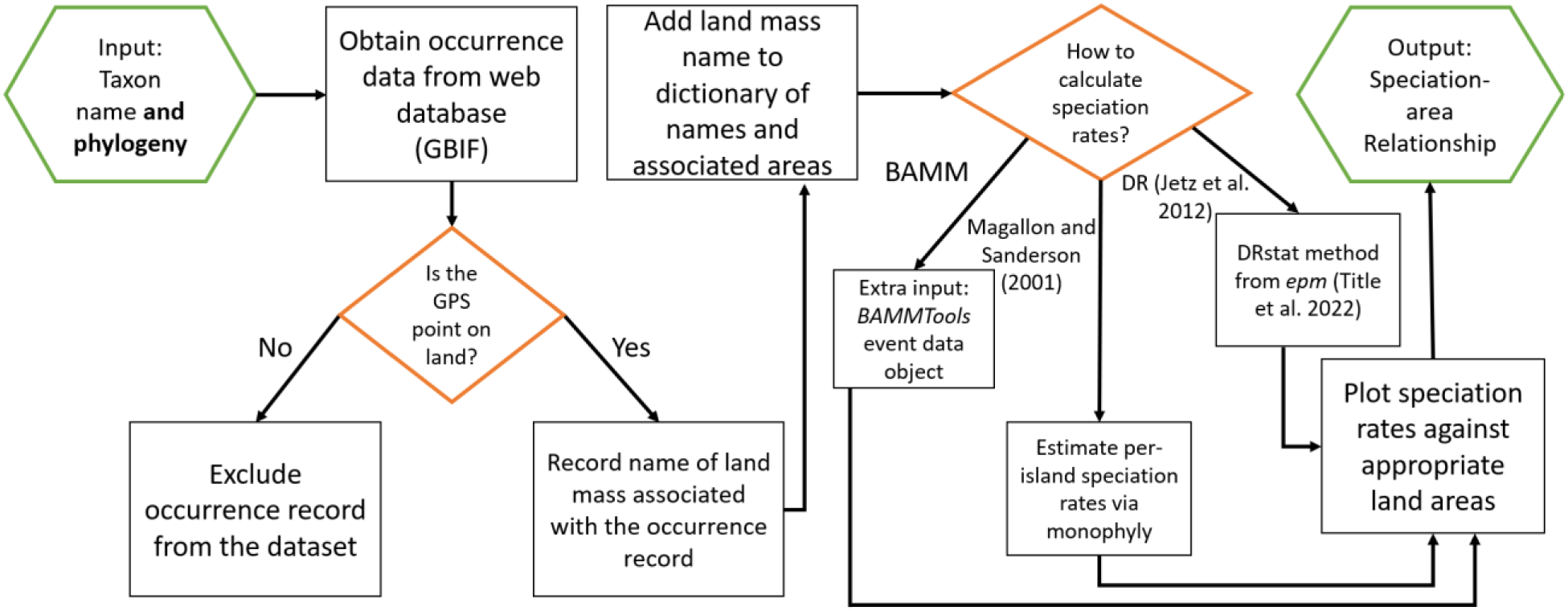
A flowchart representing the basic workflow for using *SSARP* to create a speciation-area relationship (SpAR).

**Table 2.** Functions included in *SSARP* for creating a speciation-area relationship (SpAR).

| Function | Description | Input |
| --- | --- | --- |
| speciationBAMM | Use a bammdata object created using *BAMMtools* (Rabosky et al. 2014) from a BAMM (Rabosky 2014) analysis to estimate tip speciation rates. These speciation rates are then returned with the corresponding occurrence records from previous *SSARP* workflow steps. | *BAMMtools* eventdata object, specification of the tree tip label format (binomial or species epithet), occurrence records from “findAreas” |
| speciationDR | Calculate the DR statistic (Jetz et al. 2012) using the *epm* package (Title et al. 2022) to estimate tip speciation rates. These speciation rates are then returned with the corresponding occurrence records from previous *SSARP* workflow steps. | Phylogenetic tree corresponding with taxa to be used for the SpAR, specification of the tree tip label format (binomial or species epithet), occurrence records from “findAreas” |
| speciationMS | Use methodology from Magallón and Sanderson (2001) to estimate per-island speciation rates. These speciation rates are then returned with the corresponding island areas. | Phylogenetic tree corresponding with taxa to be used for the SpAR, specification of the tree tip label format (binomial or species epithet), occurrence records from “findAreas” |
| SpeARP | Use linear or segmented regression up to a specified number of breakpoints to create speciation-area relationship plots (SpeARP). The best model as determined by AIC is returned. | Occurrence records with speciation rates from “speciationBAMM,” “speciationDR,” or “speciationMS;” maximum number of breakpoints to include in regression model selection; whether the user chose the “speciationMS” function to calculate speciation rates (TRUE) or not (FALSE) |

## Applied Example

As an example of the *SSARP* workflow, we will gather the first 10000 records from GBIF for the lizard genus *Anolis* and determine whether the *SSARP* workflow creates a species-area relationship (SAR) and a speciation-area relationship (SpAR) for island-dwelling anole lizards in the Caribbean that are comparable to the relationships presented by Losos and Schluter (2000). We do not expect the plots created in this example to exactly match the comparison from Losos and Schluter (2000) because the occurrence records are not equivalent between their paper and this example. However, the expectation is that the segmented regressions created using *SSARP* will be reasonably similar to those in Losos and Schluter (2000).

### Creating a Species-Area Relationship

*SSARP* must be installed using the “install_github” function in *devtools* (Wickham et al. 2022).

> install_github(“kmartinet/SSARP”)

> library(SSARP)

Once *SSARP* is installed and loaded, data will be collected from GBIF. While this example focuses on data from GBIF, users are able to supply their own data while executing *SSARP*’s functions to create SARs and SpARs. In order to access data from GBIF using the *rgbif* package (Chamberlain et al. 2024), the key associated with the taxon of interest in the GBIF database must be determined. This key will be found using the “getKey” function in *SSARP*:

> key <-getKey(query = “Anolis”, rank = “genus”)

> # Parameters include the taxon of interest and its rank

The “getData” function in SSARP uses GBIF’s API to access the records associated with the key obtained above. The “getData” function uses the “occ_search” function from *rgbif* (Chamberlain et al. 2024) to only return occurrence data that includes GPS coordinates. The “limit” parameter will be set to 10000 in this case for a quick illustration of *SSARP*’s functionality, but this parameter can be as large as 100,000 (the hard limit from *rgbif* for the number of records returned). If the user wants to create a SAR for a taxon above species rank that has more than 100,000 records on GBIF (the genus *Anolis*, for example, has over 300,000 georeferenced records), we recommend using individual species as queries and combining these smaller datasets to create a dataset that encompasses all possible records. This methodology is described in the Supplementary Material.

We are only interested in occurrence records for island-dwelling anole lizards located in the Caribbean, so we will geographically restrict the returned data to this area by setting the “geometry” parameter to a polygon in Well Known Text (WKT) format that encompasses the Caribbean islands.

> dat <-getData(key = key, limit = 10000, geometry = ’POLYGON((-84.8 23.9, -84.7 16.4, -65.2 13.9, -63.1 11.0, -56.9 15.5, -60.5 21.9, -79.3 27.8, -79.8 24.8, -84.8 23.9))’)

Once the occurrence data is returned, we will use each occurrence record’s GPS point to determine the land mass on which the species was found and find the area associated with that land mass using a database of island areas and names from *SSARP*.

> land <-findLand(occurrences = dat) # Finds land mass names

> area_dat <-findAreas(occs = land) # Finds land areas (in m^2^)

The “removeContinents” function in *SSARP* removes any continental occurrence records, which is useful when the user is only interested in island-dwelling species (as we are in this example).

While the data obtained by using the “getData” function was geographically restricted, potential user error in specifying the polygon in WKT format often leads to accidental continental records that will be removed by using this function.

> nocont_dat <-removeContinents(occs = area_dat)

Next, we will generate the SAR using the “SARP” function. The “SARP” function creates multiple regression objects with breakpoints up to the user-specified “npsi” parameter. For example, if “npsi” is two, “SARP” will generate regression objects with zero (linear regression), one, and two breakpoints. The function will then return the regression object with the lowest AIC score. The species-area relationship for *Anolis* in Losos and Schluter (2000) is represented by a segmented regression with one breakpoint, so “npsi” will be set to one in this example. Note that if linear regression (zero breakpoints) is better-supported than segmented regression with one breakpoint, the linear regression will be returned instead.

> SARP(occurrences = nocont_dat, npsi = 1)

The “SARP” function plots the SAR and returns a summary of the regression model. The returned SAR for island-dwelling *Anolis* data within the first 10000 records for the genus in GBIF constrained to a polygon around the Caribbean is presented in Figure 3.

**Figure 3.**
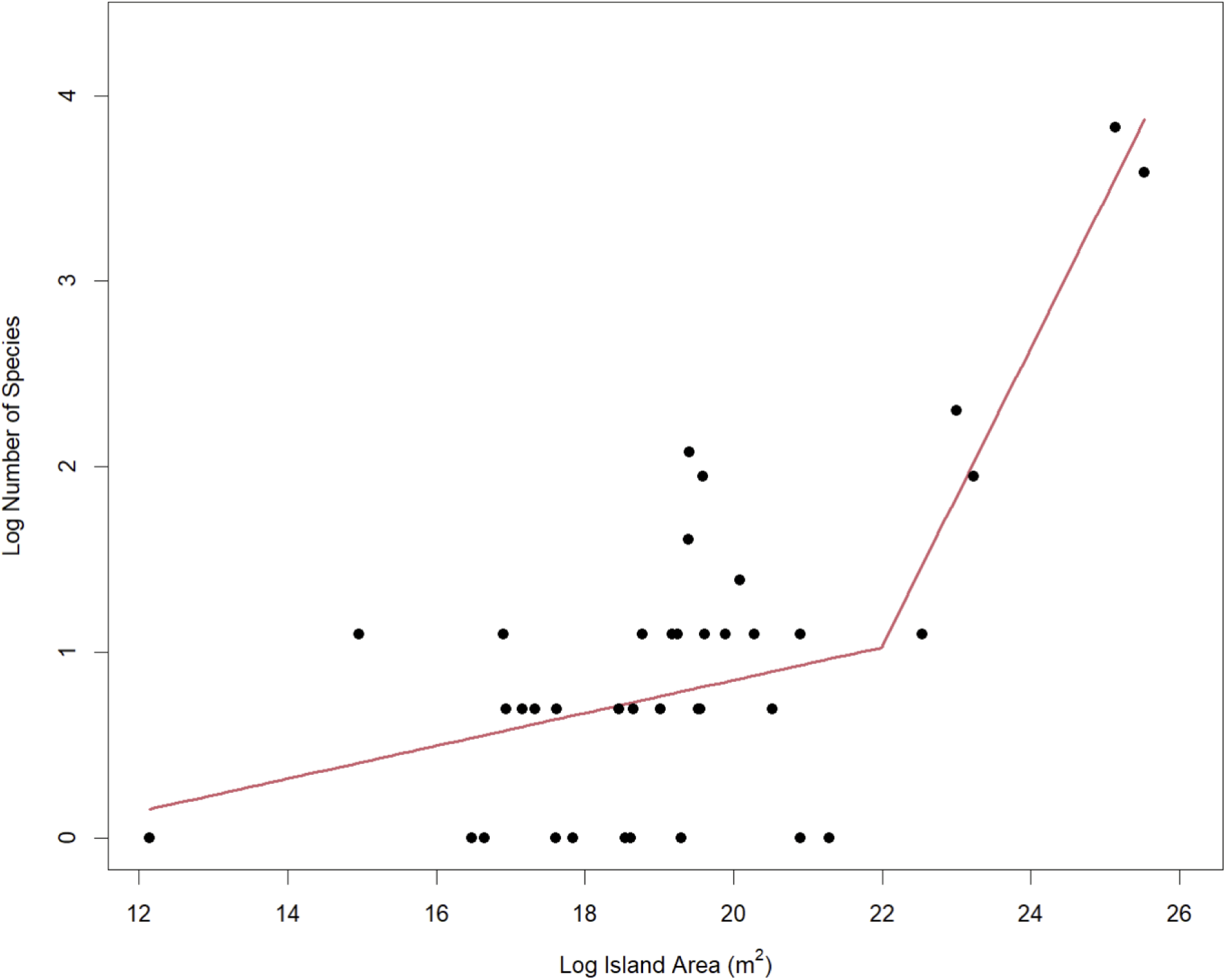
The SAR for the lizard genus *Anolis* returned by *SSARP* for the island-based occurrences within a polygon around Caribbean islands from the first 10000 records for the genus in GBIF. The estimated slopes for the SAR returned by SSARP were 0.09 and 0.71, with a breakpoint of approximately 22 log(m^2^). As a comparison, the estimated slopes for the *Anolis* SAR reported by Losos and Schluter (2000) were 0.06 and 0.76, with a breakpoint of approximately 22 log(m^2^).

This example walked through the *SSARP* workflow sequentially, but if a user would prefer to create a SAR using occurrence records from GBIF using only one command, the “quickSARP” function can be used to produce the same result as presented in Figure 3. The parameters for this function are analogous to the parameters used in the workflow above, except for the “continent” parameter that is set to “TRUE” when the user would like to remove continents from the dataset used to create the SAR.

> quickSARP(taxon = "Anolis", rank = "genus", limit = 10000,

> geometry = ’POLYGON((-84.8 23.9, -84.7 16.4, -65.2 13.9, -63.1

> 11.0, -56.9 15.5, -60.5 21.9, -79.3 27.8, -79.8 24.8, -84.8

> 23.9))’, continent = TRUE, npsi = 1)

### Creating a Speciation-Area Relationship

The “nocont_dat” object created above to generate the SAR in Figure 3 can be used with a phylogenetic tree to create a SpAR. This step in the *SSARP* workflow enables the user to determine whether the breakpoint in the SAR corresponds with a threshold for island size at which *in situ* speciation occurs (see Losos and Schluter 2000). The phylogenetic tree for *Anolis* used by Patton et al. (2021) was trimmed to only include anoles found on islands in the Caribbean for use in this example. This trimmed tree is available in *SSARP*’s GitHub repository. The SpAR presented in Losos and Schluter (2000) was generated using a speciation rate estimation method that is similar to Equation 4 in Magallόn and Sanderson (2001), so we will use the “speciationMS” function from *SSARP* to estimate speciation rates in this example. The “label_type” parameter in the “speciationMS” function corresponds to the way the tip labels are written in the user-provided phylogenetic tree. If the tip labels are simply the species epithet, as they are in the example tree here, the “label_type” parameter should be set to “epithet.” If the tip labels include the full species name, the “label_type” parameter should be set to “binomial.”

> tree <-read.tree(file = "Patton_Anolis_Trimmed.tree") # Read in tree

> speciation_occurrences <-speciationMS(tree = tree, label_type =

> “epithet”, occurrences = nocont_dat) # Calculate speciation rates

The newly created “speciation_occurrences” object is a dataframe containing island areas with their corresponding speciation rate as estimated by the “speciationMS” function. Next, we will use the “speciation_occurrences” object with the “SpeARP” function to create a SpAR. We will again set the “npsi” parameter to one because the SpAR presented in Losos and Schluter (2000) has one breakpoint. Note that if linear regression (zero breakpoints) is better-supported than segmented regression with one breakpoint, the linear regression will be returned instead. The final parameter is the “MS” parameter, which tells the function whether the user generated speciation rate estimates using the “speciationMS” function (TRUE) or not (FALSE). This is an important parameter because if “speciationMS” is used, the speciation rate calculation already log-transforms the values. The other speciation rate estimation methods do not automatically log-transform the speciation rate values.

> SpeARP(occurrences = speciation_occurrences, npsi = 1, MS = TRUE)

The “SpeARP” function plots the SpAR and returns a summary of the regression model. The returned speciation-area for the same island-dwelling *Anolis* data we gathered for the SAR is presented in Figure 4.

**Figure 4.**
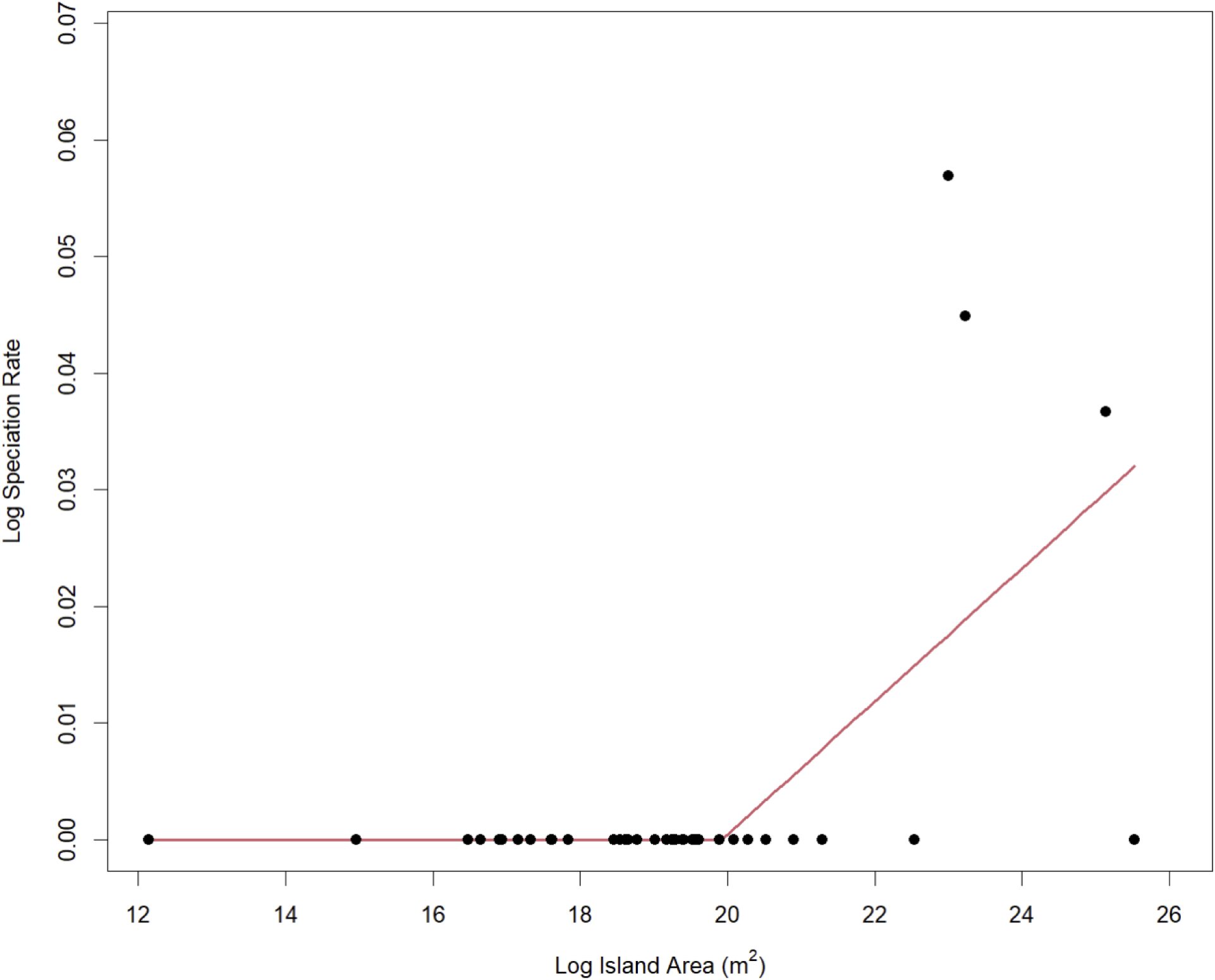
The speciation-area relationship (SpAR) for the lizard genus *Anolis* returned by *SSARP* for the island-based occurrences within a polygon around Caribbean islands from the first 10000 records for the genus in GBIF. The estimated breakpoint for the SpAR returned by *SSARP* was 19.9 log(m^2^). The breakpoint for the *Anolis* SpAR reported by Losos and Schluter (2000) was approximately 22 log(m^2^). Unlike the results from Losos and Schluter (2000), the breakpoint estimated by *SSARP* for this SpAR does not match the SAR breakpoint. This very likely occurred because the calculation for speciation rate in Magallόn and Sanderson (2001) that “speciationMS” uses is based on monophyly, which can be disrupted on islands with non-native species occurrence records.

## Methods

### Finding Island Areas

Part of the workflow for *SSARP* is to determine whether an occurrence record’s GPS point truly corresponds with a land mass. The “findLand” function uses the “map.where” function in the *maps* R package (Becker et al. 2023), which returns the name of the land mass associated with a GPS point input. Multiple databases are tested in this process to attempt to fill in any gaps left over from each database reference. First, the “worldHires” database from the *mapdata* R package (Becker et al. 2022) is used with the “map.where” function. Next, the “world” database from *mapdata* is used to attempt to fill in any gaps left over from using the “worldHires” database. Finally, if the “fillgaps” argument in the “findLand” function is set to “TRUE,” the Photon API (komoot 2024) will be queried for each GPS point that did not receive a land mass name from the “map.where” calls. Photon provides an easy method of accessing the OpenStreetMap API (OpenStreetMap contributors 2024) and returns detailed information about the location associated with a GPS point. The information useful for creating SARs for island species in *SSARP*, such as country and island name, is sometimes listed in different parts of the data returned by Photon. Considering the structure of the Photon output, “findLand” saves three sections of the Photon result: country, locality, and county. These three parameters were found to most reliably include the country and island names for a wide variety of GPS points associated with islands across the globe.

One of the most important components of *SSARP* is a dataset of island names and their associated areas. This dataset was created using *ArcPy*, a Python library for conducting geographic analyses with ArcGIS Pro (ESRI 2024). The scripts used to gather all of the island data is accessible in *SSARP*’s GitHub repository (Martinet 2024). Global island data from Sayre et al. (2018) was queried from the “Default” geodatabase in ArcGIS Pro using three separate environment masks: one for islands with an area smaller than 1 km^2^, one for islands with an area larger than 1 km^2^, and one for continents. The elevation of each island was also recorded. The “ZonalStatisticsAsTable” function was used to compile the spatial data and output it as a csv file for use in *SSARP*. Island areas were approximated by ArcGIS Pro through the use of pixel counts. Each pixel represented a 250 m x 250 m (62500 m^2^) area, and the reported area for each land mass was calculated by multiplying the number of pixels that cover a land mass by the area of one pixel.

### Speciation Rate Estimation

Three methods for calculating speciation rates are included in *SSARP*: BAMM (Rabosky 2014), DR (Jetz et al. 2012), and the lambda calculation from Magallόn and Sanderson (2001). While tip speciation rate estimations, as calculated in BAMM and DR, are not useful for every analysis, examining the speciation-area relationship (SpAR) for taxa is a good use case for tip speciation rates because these relationships focus on non-historical geographic patterns of diversity (Title and Rabosky 2019). The “speciationBAMM” function in *SSARP* requires a bammdata object as input, which must be created using the *BAMMtools* package (Rabosky et al. 2014) after the user completes a BAMM analysis. This object includes tip speciation rates by default in the “meanTipLambda” list element, which *SSARP* accesses to add the appropriate tip speciation rates for each species to the occurrence record dataframe.

DR stands for “diversification rate,” but it is ultimately a better estimation of speciation rate than net diversification (Belmaker and Jetz 2015; Quintero and Jetz 2018) and returns results similar to BAMM’s tip speciation rate estimations (Title and Rabosky 2019). Due to the nature of this metric, the “speciationDR” function returns the values obtained from running the “DRstat” function from the *epm* package (Title et al. 2022) as tip speciation rates.

In addition to tip speciation rates, *SSARP* includes a function for calculating the speciation rate for a clade from Magallόn and Sanderson (2001). The “speciationMS” function in *SSARP* uses the “subtrees” function from *ape* (Paradis and Schliep 2019) to generate all possible subtrees from the user-provided phylogenetic tree that corresponds with the taxa of interest for the SpAR. Then, species in the provided occurrence records generated from previous steps in the *SSARP* workflow are grouped by island. For each group of species that comprise an island, the number of subtrees that represent that group of species and the root age of each subtree is recorded, along with the name and area of the island. The speciation rate for each subtree is then calculated following Equation 4 in Magallόn and Sanderson (2001). If an island includes multiple subtrees, the island speciation rate is the average of the calculated speciation rates. This average is calculated when the SpAR is plotted. When the “SpeARP” function from *SSARP* is used to plot the SpAR, the user must specify whether “speciationMS” was used to calculate speciation rates. This distinction is important because the Magallόn and Sanderson (2001) method already log-transforms the value for speciation rate, and plotting with “SpeARP” would log-transform these rates again if the user does not specify whether “speciationMS” was used.

### Caveats

Given the nature of data from online databases such as GBIF, occurrence records used for creating SARs using *SSARP* might need to be filtered more rigorously than the filtering mechanisms already included in the *SSARP* workflow. For example, the user might want to remove occurrence records that correspond with non-native observations of the taxon of interest because these records might skew the resulting SAR (Baiser and Li 2018; Guo et al. 2021). These records would similarly skew resulting SpARs, especially when using the “speciationMS” function to calculate speciation rates due to the importance of clades in the equation used in that function. Additionally, if a GPS coordinate in the occurrence dataset is dramatically incorrect, a land mass that should not be included in the taxon’s range might be included in the relationship and create an outlier. These outliers are often visually obvious when the plot is created and the faulty occurrence record can be easily spotted for removal from the dataframe created by the “findAreas” function.

## Conclusions

The *SSARP* package provides users with a seamless workflow for gathering occurrence data from GBIF and creating species-area relationships (SARs) and speciation-area relationships (SpARs) with that data. Before the creation of *SSARP*, researchers who wanted to create these relationships using information from online occurrence databases such as GBIF would have needed to install several packages for gathering the data, filtering the data, and creating the plot itself. Additionally, researchers previously had to conduct extensive literature searches or trace land masses on a mapping program to assemble their own dataset of island areas pertinent to their study system in order to create a SAR or SpAR. *SSARP* precludes this previous need to install several individual packages and includes a dataset of areas for islands across the globe. While the process of creating SARs and SpARs in *SSARP* is built on powerful methods, outliers might still emerge. Researchers should examine resulting plots carefully to ensure that none of the occurrence data returned by GBIF represents land masses or taxa that should not be included in the final relationship when considering the study design. The ease with which researchers create species-area relationships and speciation-area relationships using *SSARP* will allow for the emergence of more studies that compare these relationships using global datasets, which will hopefully lead us to a clearer picture of the world’s biodiversity.

## Code Availability

The *SSARP* R package and external data used in the Applied Example are available freely on GitHub in the following repository: https://github.com/kmartinet/SSARP. We plan to submit *SSARP* to CRAN and rOpenSci.

## Acknowledgements

We thank Bruce Godfrey, GIS Librarian at the University of Idaho, for helping us generate the database of island names and associated areas used in *SSARP*. KM was supported by the University of Idaho Institute for Interdisciplinary Data Sciences (IIDS), Bioinformatics and Computational Biology Program (BCB), and Department of Biological Sciences.

## Literature Cited

Arrhenius, O. (1921). Species and Area. Journal of Ecology, 9(1): 95–99.

Baiser, B. & Li, D. (2018). Comparing species–area relationships of native and exotic species. Biological Invasions, 20: 3647–3658.

Becker, R.A., Wilks, A.R., & Brownrigg, R. (2022). mapdata: Extra Map Databases. R package version 2.3.1, https://CRAN.R-project.org/package=mapdata

Becker, R.A., Wilks, A.R., Brownrigg, R., Minka, T.P., & Deckmyn, A. (2023). maps: Draw Geographical Maps. R package version 3.4.2, https://CRAN.R-project.org/package=maps

Belmaker, J., & Jetz, W. (2015). Relative roles of ecological and energetic constraints, diversification rates and region history on global species richness gradients. Ecology Letters, 18: 563–571.

Chamberlain, S., Barve, V., Mcglinn, D., Oldoni, D., Desmet, P., Geffert, L., Ram, K. (2024). rgbif: Interface to the Global Biodiversity Information Facility API. R package version 3.7.8, https://CRAN.R-project.org/package=rgbif

Chisholm, R.A., Lim, F., Yeoh, Y.S., Seah, W.W., Condit, R., & Rosindell, J. (2018). Species-area relationships and biodiversity loss in fragmented landscapes. Ecology Letters, 21: 804–813.

de Silva, M.A.F., Mendes, C.B., & Prevedello, J.A. (2024). How important is passive sampling to explain species-area relationships? A global synthesis. Landscape Ecology, 39: 50.

ESRI. (2024). ArcPy, Python library, https://developers.arcgis.com/documentation/arcgis-add-ins-and-automation/arcpy/

Fasola S., Muggeo V.M.R., Kuchenhoff K. (2018). A heuristic, iterative algorithm for change-point detection in abrupt change models. Computational Statistics, 33: 997–1015.

Gleason, H.A. (1922). On the Relation Between Species and Area. Ecology, 3(2): 158–162.

Guo, Q., Cen, X., Song, R., McKinney, M.L., Wang, D. (2021). Worldwide effects of non-native species on species-area relationships. Conservation Biology, 35(2): 711–721.

Heaney, L.R. (2000). Dynamic disequilibrium: A long-term, large-scale perspective on the equilibrium model of island biogeography. Global Ecology and Biogeography, 9(1): 59– 74.

Jetz, W., Thomas, G.H., Joy, J.B., Hartmann, K., & Mooers, A.O. (2012). The global diversity of birds in space and time. Nature, 491: 444–448.

komoot. (2024). Photon API. Retrieved from https://photon.komoot.io/

Losos, J.B. & Schluter, D. (2000). Analysis of an evolutionary species-area relationship. Nature, 408: 847–850.

MacArthur, R.H. & Wilson, E.O. (1967). The Theory of Island Biogeography. Princeton, N.J: Princeton University Press.

Magallόn, S. & Sanderson, M.J. (2001). Absolute Diversification Rates in Angiosperm Clades. Evolution, 55(9): 1762–1780.

Martinet, K.M. (2024). SSARP: Species-/Speciation-Area Relationship Projector. R package version 0.0.1. https://github.com/kmartinet/SSARP

Matthews, T.J., Triantis, K.A., Whittaker, R.J., & Guilhaumon, F. (2019). sars: an R package for fitting, evaluating and comparing species–area relationship models. Ecography, 42: 1446–1455.

Matthews, T.J. & Rigal, F. (2021). Thresholds and the species-area relationship: a set of functions for fitting, evaluating and plotting a range of commonly used piecewise models in R. Frontiers of Biogeography, 13(1): e49404.

OpenStreetMap contributors. (2024). Retrieved from https://planet.openstreetmap.org

Owens, H., Barve, V., Chamberlain, S. (2024). spocc: Interface to Species Occurrence Data Sources. R package version 1.2.3, https://CRAN.R-project.org/package=spocc

Paradis, E. & Schliep, K. (2019). ape 5.0: an environment for modern phylogenetics and evolutionary analyses in R. Bioinformatics, 35: 526–528.

Patton, A.H., Harmon, L.J., del Rosario Castañeda, M., Frank, H.K., Donihue, C.M., Herrel, A., & Losos, J.B. (2021). When adaptive radiations collide: Different evolutionary trajectories between and within island and mainland lizard clades. PNAS, 118(42): e2024451118.

Quintero, I., & Jetz, W. (2018). Global elevational diversity and diversification of birds. Nature, 555, 246–250.

Rabosky, D.L. (2014). Automatic Detection of Key Innovations, Rate Shifts, and Diversity-Dependence on Phylogenetic Trees. PLOS ONE, 9(2): e89543.

Rabosky, D.L., Grundler, M., Anderson, C., Title, P., Shi, J.J., Brown, J.W., Huang, H., & Larson, J.G. (2014), BAMMtools: an R package for the analysis of evolutionary dynamics on phylogenetic trees. Methods in Ecology and Evolution, 5: 701–707.

Rosindell, J., & Harmon, L.J. (2013). A unified model of species immigration, extinction and abundance on islands. Journal of Biogeography, 40(6): 1107–1118.

Rosenzweig, M.L. (1995). Species Diversity in Space and Time. Cambridge, England: Cambridge University Press.

Scheiner, S.M. (2003). Six types of species-area curves. Global Ecology & Biogeography, 12: 441–447.

Sayre, R., Noble, S., Hamann, S., Smith, R., Wright, D., Breyer, S., Butler, K., Van Graafeiland, K., Frye, C., Karagulle, D., Hopkins, D., Stephens, D., Kelly, K., Basher, Z., Burton, D., Cress, J., Atkins, K., Van Sistine, D.P., Friesen, B., Allee, B., Allen, T., Aniello, P., Asaad, I., Costello, M.J., Goodin, K., Harris, P., Kavanaugh, M., Lillis, H., Manca, E., Muller-Karger, F., Nyberg, B., Parsons, R., Saarinen, J., Steiner, J., & Reed, A. (2018). A new 30 meter resolution global shoreline vector and associated global islands database for the development of standardized global ecological coastal units. Journal of Operational Oceanography, 12(S2): S47–S56. 10.1080/1755876X.2018.1529714

Title, P., Swiderski, D., & Zelditch, M. (2022). EcoPhyloMapper: an R package for integrating geographic ranges, phylogeny, and morphology. Methods in Ecology and Evolution, 13: 1912–1922.

Wagner, C.E., Harmon, L.J., & Seehausen, O. (2014). Cichlid species-area relationships are shaped by adaptive radiations that scale with area. Ecology Letters, 17(5): 583–592.

Whittaker, R.J., Triantis, K.A., & Ladle, R.J. (2008). A general dynamic theory of oceanic island biogeography. Journal of Biogeography, 35(6): 977–994.

Wickham, H., Hester, J., Chang, W., Bryan, J. (2022). devtools: Tools to Make Developing R Packages Easier. R package version 2.4.5, https://CRAN.R-project.org/package=devtools

Zizka, A., Silvestro, D., Andermann, T., Azevedo, J., Duarte Ritter, C., Edler, D., Farooq, H., Herdean, A., Ariza, M., Scharn, R., Svanteson, S., Wengtrom, N., Zizka, V. & Antonelli, A. (2019). CoordinateCleaner: standardized cleaning of occurrence records from biological collection databases. Methods in Ecology and Evolution, 10(5):744–751.

