## Supplemental Text for "*SSARP*: An R Package for Easily Creating Species- and Speciation- Area Relationships Using Web Databases"

### Creating Large Occurrence Record Datasets

The "getData" function in *SSARP* uses the "occ_search" function from *rgbif* (Chamberlain et al. 2024) to only return occurrence data that includes GPS coordinates. In the Worked Example in the main text, the "limit" parameter was set to 10000 for a quick illustration of *SSARP*'s functionality. This parameter can be as large as 100,000, which is the hard limit from *rgbif* for the number of records returned. If the user wants to create a SAR for a taxon above species rank that has more than 100,000 records on GBIF (the genus *Anolis*, for example, has over 300,000 georeferenced records), we recommend using individual species as queries and combining these smaller datasets to create a dataset that encompasses all possible records.

A user who wants to create a large dataset from multiple species-level queries using can write a loop in R that uses the "getKey" and "getData" functions from *SSARP* with each species, and then combine the dataframes returned from the "getData" function. The code chunk below provides an example of this methodology:

```
# Five random anoles

species_list <- c("Anolis allogus", "Anolis rubribarbus",
"Anolis equestris", "Anolis vermiculatus", "Anolis
alutaceus")
```

```
# Create empty dataframe
occ_df <- data.frame()


# Loop over species in species list
for (sp in species_list){
  # Get key for current species
  key <- getKey(query = sp, rank = "species")
  # Get data for current species, maximum limit
  dat <- getData(key = key, limit = 100000)


  # Sometimes rgbif returns dataframes of different sizes
with different column names
  # Select the most important columns:
  sp_df <- dat[c("scientificName", "acceptedScientificName",
"decimalLatitude", "decimalLongitude", "basisOfRecord",
"occurrenceStatus", "genericName", "specificEpithet",
"continent", "stateProvince", "references", "license",
"rightsHolder")]


  # Add information from current species to large dataframe
```

```
  occ_df <- rbind(occ_df, sp_df)
}
# occ_df now includes all occurrence records from GBIF for
all of the specified anoles
```

**Literature Cited**

Chamberlain, S., Barve, V., Mcglinn, D., Oldoni, D., Desmet, P., Geffert, L., Ram, K. (2024). rgbif: Interface to the Global Biodiversity Information Facility API. R package version 3.7.8, https://CRAN.R-project.org/package=rgbif
